# Infrared heater warming system markedly reduces dew formation: An overlooked factor in arid ecosystems

**DOI:** 10.1101/2020.02.15.950584

**Authors:** Tianjiao Feng, Lixu Zhang, Qian Chen, Zhiyuan Ma, Hao Wang, Zijian Shangguan, Lixin Wang, Jin-Sheng He

**Author notes:** Co-first authors: Tianjiao Feng and Lixu Zhang have equal contribution to this study. Corresponding author: Jin-Sheng He: State Key laboratory of Grassland Agro-ecosystems, College of Pastoral Agriculture Science and Technology, Lanzhou University, Lanzhou 730020, China.

## Abstract

Dew plays a vital role in ecosystem processes in arid and semi-arid regions and is expected to be affected by climate warming. Infrared heater warming systems have been widely used to simulate climate warming effects on ecosystem. However, how this warming system affects dew formation has been long ignored and rarely addressed. In a typical alpine grassland ecosystem on the Northeast of the Tibetan Plateau, we measured dew amount and duration by artificial condensing surfaces, leaf wetness sensors and *in situ* dew formation on plants from 2012 to 2017. We also measured plant traits related to dew conditions. The results showed that (1) warming reduced the dew amount by 41.6%-91.1% depending on the measurement method, and reduced dew duration by 32.1 days compared to the ambient condition. (2) Different plant functional groups differed in dew formation. (3) Under the infrared warming treatment, the dew amount decreased with plant height, while under the ambient conditions, the dew amount showed the opposite trend. We concluded that warming with an infrared heater system greatly reduces dew formation, and if ignored, it may lead to overestimation of the effects of climate warming on ecosystem processes in climate change simulation studies.

## 1. Introduction

Dew is considered a vital vegetative water source in semiarid and arid areas (Beysens, 1995; Agam and Berliner, 2006; Wang et al., 2017a). In such environments, dew plays an indispensable role on plants (Benasher et al., 2010; Zhuang and Ratcliffe, 2012; Oliveira, 2013), biological crusts (Zhang et al., 2009; Fischer et al., 2012; Kidron and Temina, 2013), small animals (Steinberger et al., 1989; Zheng et al., 2010) and plant-associated microorganisms (Agam and Berliner, 2006). Dew also has significant effects on relative humidity, vapor pressure deficits and nutrient cycling (Munné, 1999; Goldsmith et al., 2013; Wang et al., 2019), and these factors influence plant photosynthesis, transpiration and other important ecological processes (Benasher et al., 2010; Wang et al., 2017a).

Dew formation in ecosystems is affected by microclimatic parameters (e.g., air temperature, relative humidity and vapor pressure deficit) and plant morphological features (e.g., aboveground biomass, leaf area, leaf roughness and plant height). These factors change under different climatic conditions and are associated with different plant species or functional groups (Agam and Berliner, 2006; Hao et al., 2012). Thus, it is expected that rapidly changing climates will significantly affect dew formation (Walther et al., 2002; Xiao et al., 2013; Li et al., 2018).

To simulate climate warming, an infrared heater warming system is widely used to address the potential impacts of climate warming on ecosystems in the field (Liu et al., 2018; Song et al., 2019; Ettinger et al., 2019). However, there are differences between artificial warming and natural warming (Shaver et al., 2000) and the effects of artificial warming have the potential to influence dew formation (Wolkovich et al., 2012; Moni et al., 2019).

Recently, there are increasing number of studies on dew research (Tomaszkiewicz et al., 2015; Wang et al., 2017a), most of which analyzed the ecological effects of dew on ecosystem processes, such as plant photosynthesis and transpiration in ecosystems (Ninari and Berliner, 2002; De Boeck et al., 2015; Wang et al., 2019), or compared the effects of environmental factors on dew formation (Hao et al., 2012; Ettinger et al., 2019; Beysens, 2016). There are also substantial efforts have been made to study the potential impacts of climate warming on dryland ecosystems by manipulating temperature in the field with various warming facilities (Kimball et al., 2018; Moni et al., 2019; Song et al., 2019). However, the effects of artificial warming on dew formation and ecosystem processes have not been addressed. As a result, the observed changes in ecological processes in various climate change studies are likely attributed, to some extent, to altered dew amounts, misrepresenting the effects of warming on ecosystem processes.

Few studies on dew research have been conducted in the context of climate change, and global warming experiments have not reported the effects of climate change or plant traits and functional groups on dew formation or even considered the effects of dew as a long-term factor affecting soils and plants as well as ecosystem processes during the course of climate change (Tomaszkiewicz et al., 2015; Li et al., 2018). On the other hand, few studies have investigated the influences of different plant traits or functional groups on dew amount and duration (Wang et al., 2012; Xu et al., 2015; Tomaszkiewicz et al., 2016). Therefore, the impacts of artificial warming, plant traits and functional groups on dew formation urgently need to be revealed to better understand the impacts of warming on ecosystem processes (Korell et al., 2019).

Experimental data from field-based climate change experiments are crucial to determine mechanistic links between simulated climate change and dew formation. This study is a part of a comprehensive warming experiment in a typical alpine grassland in Tibet Plateau, where we measured the dew amount and duration by the methods of the artificial condensation surface, leaf wetness sensor and *in situ* plant dew formation measurement to explore the responses of dew formation among different functional groups to simulated climate warming. The objectives of the present study were to (1) address how the widely used infrared heater warming system affects dew amount and duration and (2) elucidate whether plant functional groups, which are expected to shift under future warming, affect dew formation under ambient and warming conditions. Our results will aid in the understanding of the characteristics of dew formation under a warming climate in the future.

## 2. Materials and methods

### 2.1. Study site

The study site was located at Haibei National Field Research Station of the Alpine Grassland Ecosystem (37°36′ N, 101°19′ E, 3215 m a.s.l.) in the northeastern part of the Tibetan Plateau, China. The mean annual air temperature and precipitation were −1.2 °C and 489.0 mm during 1980-2014, respectively (Liu et al., 2018). Approximately 80% of the precipitation was concentrated in the growing season (from May to September). This mesic alpine grassland is dominated by *Stipa aliena, Elymus nutans* and *Helictotrichon tibeticum*. The soil is classified as a Mollisol according to USDA Soil Taxonomy. The average soil bulk density, organic carbon concentration and pH were 0.8 g·cm^−3^, 63.1 g·kg^-1^ and 7.8 at the 0-10 cm soil depth, respectively (Lin et al., 2016).

### 2.2. Warming experiment design

Our study was conducted within an experimental warming × precipitation infrastructure within an area of 50 m × 110 m that was established in July 2011. The design of the experiment was detailed in Liu et al. (2018). In brief, the experiment manipulated the temperature (+2 °C, control) and precipitation (+50%, control, −50%) with a completely randomized design. Each treatment had six replications with a plot area of 2.2 m × 1.8 m (Liu et al., 2018). The warming treatment was applied by two infrared heaters (220 V, 1200 W, 1.0 m long, and 0.22 m wide) (Ma et al., 2017). In the current study, we only compared ambient and warming conditions.

Air temperature and relative humidity probes (VP-3, METER Group, Inc., Pullman, WA, USA) were installed 30 cm above the soil surface within each plot. All data were automatically recorded hourly and stored in a data logger (EM50, METER Group, Inc., Pullman, WA, USA).

### 2.3. Dew formation measurements

We used three methods to measure dew amount and duration:

#### (1) Artificial condensation surface

The daily dew production was collected and measured using a preplaced plastic film, 20 cm × 20 cm in size, at each plot (Vuollekoski et al., 2015). The clean plastic films were weighed and placed at each plot at 20:00 pm (local time) the day before each measurement. At 6:00 am the next morning, the preplaced plastic films were weighed, and the differences in the weights were designated as the dew production (g) for that night. The dew amount (mm) was equal to the dew weight divided by the area of the plastic film. In this study, the dew amounts were measured by this method on sunny and windless days two times per week during the peak growing seasons (from July to September) in 2012 and 2013.

#### (2) *In situ* dew formation measurements on plants

Dew formation on plants was measured by sampling the outside plots to avoid disturbing the plant community composition of each plot. Similar individuals of the same species were chosen to measure dew formation. For each species, four or five individuals were selected, weighed, measured plant heights and placed into floral foam to prevent wilting the day before measurement and then placed at each plot at 20:00 pm (local time). At 6:00 am the next morning, these plants were weighed after being brought back to the laboratory to attain the total weight. The dew production (g) was equal to the total weight minus the plant fresh weight. At the same time, we scanned the leaf area of plants and finally calculated the dew amount (mm) produced per unit plant area. In this study, the dew amounts were measured by this method on sunny and windless days three times per week during the peak growing season (from July to September) in 2017.

#### (3) Leaf wetness sensors

The dew amount and duration were monitored hourly using leaf wetness sensors (S-LWA-M003, Onset Computer Corporation, Bourne, MA, USA) and a HOBO data logger (H21-002, Onset Computer Corporation, Bourne, MA, USA) at each plot from 2015 to 2017 (Chen 2015). The dew amount was calculated by the fitting relationship between the measured leaf wetness sensor readings and the actual condensed water amount (g). We sprayed water evenly on the leaf wetness sensors to induce water condensation on their surface, recorded the instrument reading, and established the relationship between the condensation amount and the leaf wetness sensor readings. In addition, the simulated solid condensation amount was determined using the same method in a −20 °C refrigerator to establish a relationship curve. We repeated the above steps multiple times to ensure a wide range of leaf wetness sensor readings. The relationship curve between the leaf wetness sensor readings and the condensation amount was fitted (Fig. S1), and the relationship was as follows:

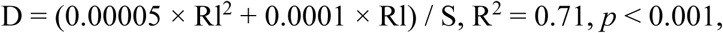

where *D* is the dew amount (mm), *Rl* is the leaf wetness sensor reading and *S* is the area of the leaf wetness sensor, which was 4.7 cm × 5.1 cm.

In our study, the former two measurement methods focused on dew amount, while only the leaf wetness sensor method measured the dew duration. The data were automatically recorded hourly, and dew duration was calculated as the number of days for which dew was recorded between 8:00 p.m. and 6:00 a.m. of the next morning during the measuring periods.

### 2.4. Dew formation and aboveground biomass at the species level

In total, we measured dew formation at the species level for 10 species. These ten species accounted for approximately 72% of the total community biomass (Liu et al., 2018). We divided these plant species into three functional groups, i.e., grasses (*Stipa aliena, Elymus nutans* and *Helictotrichon tibeticum*), forbs (*Tibetia himalaica, Oxytropis ochrocephala, Medicago ruthenica, Gentiana straminea* and *Saussurea pulchra*), and sedges (*Kobresia humilis* and *Carex przewalskii*) and separately analyzed their dew formation responses to warming. The aboveground biomass was separated into grasses, sedges, and forbs, harvested and oven-dried at 65 °C to a constant weight. Plant height was measured using five selected individuals per species in each plot before dew formation measurement during the experimental periods.

### 2.5. Data Analysis

Based on long-term meteorological observations, the dew point temperature was calculated by Penman-Monteith equation with the following function (Allen et al., 1998):

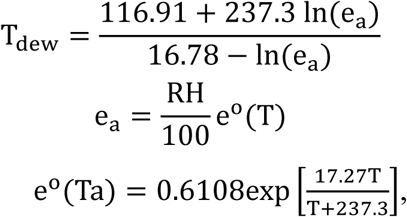

where *T*_*dew*_ is dew point temperature [°C], *e*_*a*_ is actual vapour pressure [kPa], *e*^*o*^ (T) is saturation vapor pressure at the air temperature *T*_*a*_ [kPa], and *T*_a_ is air temperature [°C]. Meanwhile, the temperature differences (T_a_-T_dew_) was calculated by the difference between the air temperature (T_a_) and dew point temperature (T_dew_) to represent the difficult degrees of dew formation.

The dew point temperature was calculated using long-term meteorological observations. Linear regression was used to test the relationship between plant height and dew amount in the control and warming treatments. To test the warming effect, one-way analysis of variance (ANOVA) and Tukey’s HSD test were used to determine differences in dew amount and duration between the control and warming plots. All statistical analyses were conducted using R 3.2.2 software (R Foundation for Statistical Computing, Vienna, Austria, 2013). Differences were considered significant at *P* < 0.05 unless otherwise stated.

## 3. Results

### 3.1. Effects of warming on the dew formation

The multiple measurement methods showed decreased dew amounts under warming conditions. Warming resulted in average decreases of 91.7%, 83.9% and 41.6% in dew amount by the artificial condensation surface method, the in situ dew formation on plants and the leaf wetness sensors, respectively (linear mixed-effects model: *P* < 0.001; Fig. 1). From 2015 to 2017, warming significantly decreased the dew duration by an average of 10.3% (linear mixed-effects model: *P* < 0.001; Fig. 2a). Therefore, warming reduced the total dew formation by not only reducing the daily dew amounts (mm/day) but also the dew duration (days). The results also showed that warming significantly increased the temperature differences (T_a_-T_dew_) by 3.8% (*P* < 0.001; Fig. 2b), which made dew formation more difficult. Furthermore, the differences in the dew amount between the control and warming treatments (D_control_-D_warming_) showed significant differences at the seasonal scale (Fig. 2c). The dew amounts under the warming treatment decreased by an average of 0.05 mm (up to 64.5%) during the growing seasons and only decreased by an average of 0.006 mm (only 27.5%) in non-growing seasons (Fig. 2c).

**Fig. 1.**
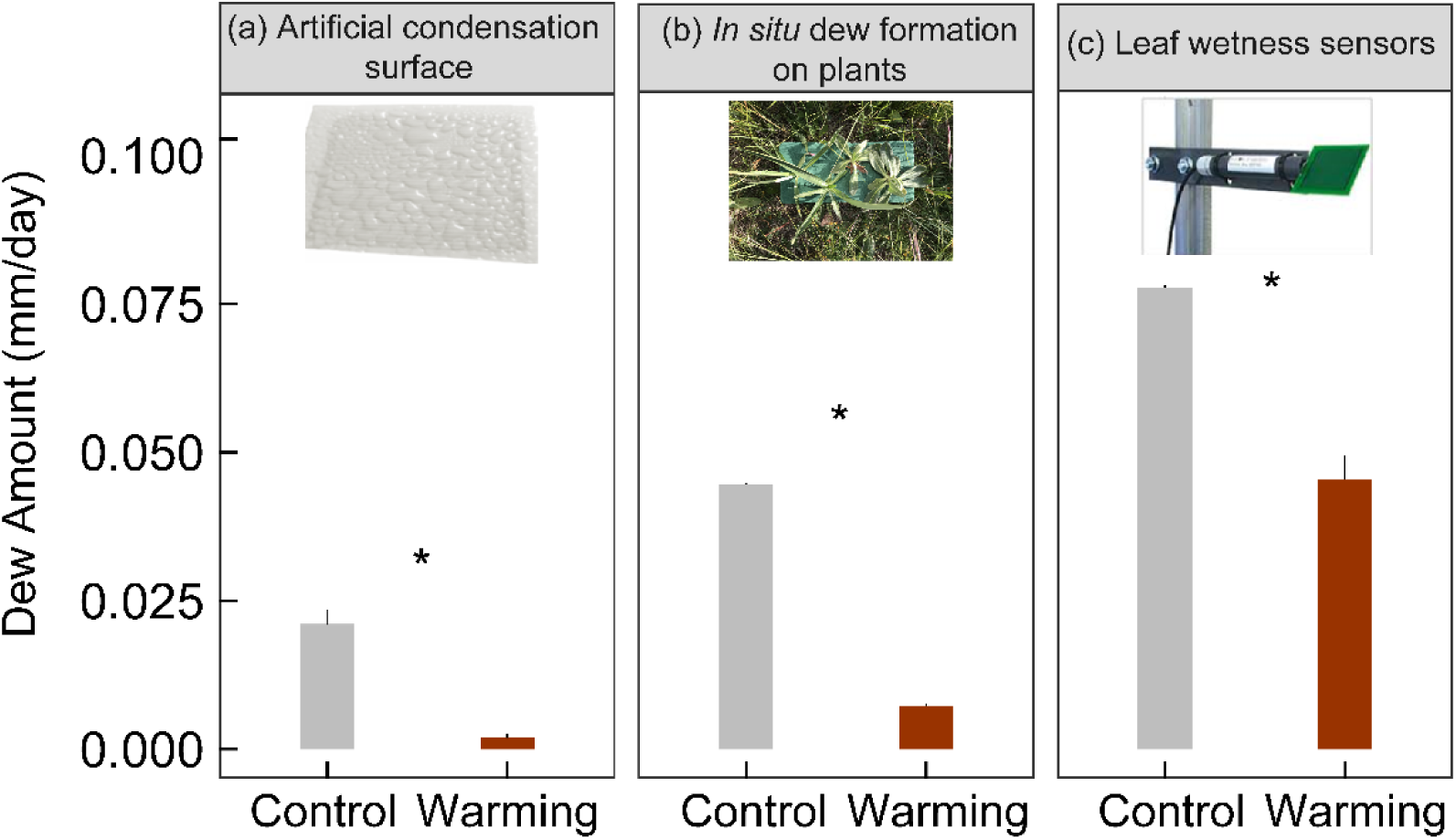
The dew amount measured by (a) artificial condensation surface, (b) *in situ* dew formation on plants and (c) leaf wetness sensors in control and warming treatments during the experimental period.

**Fig. 2.**
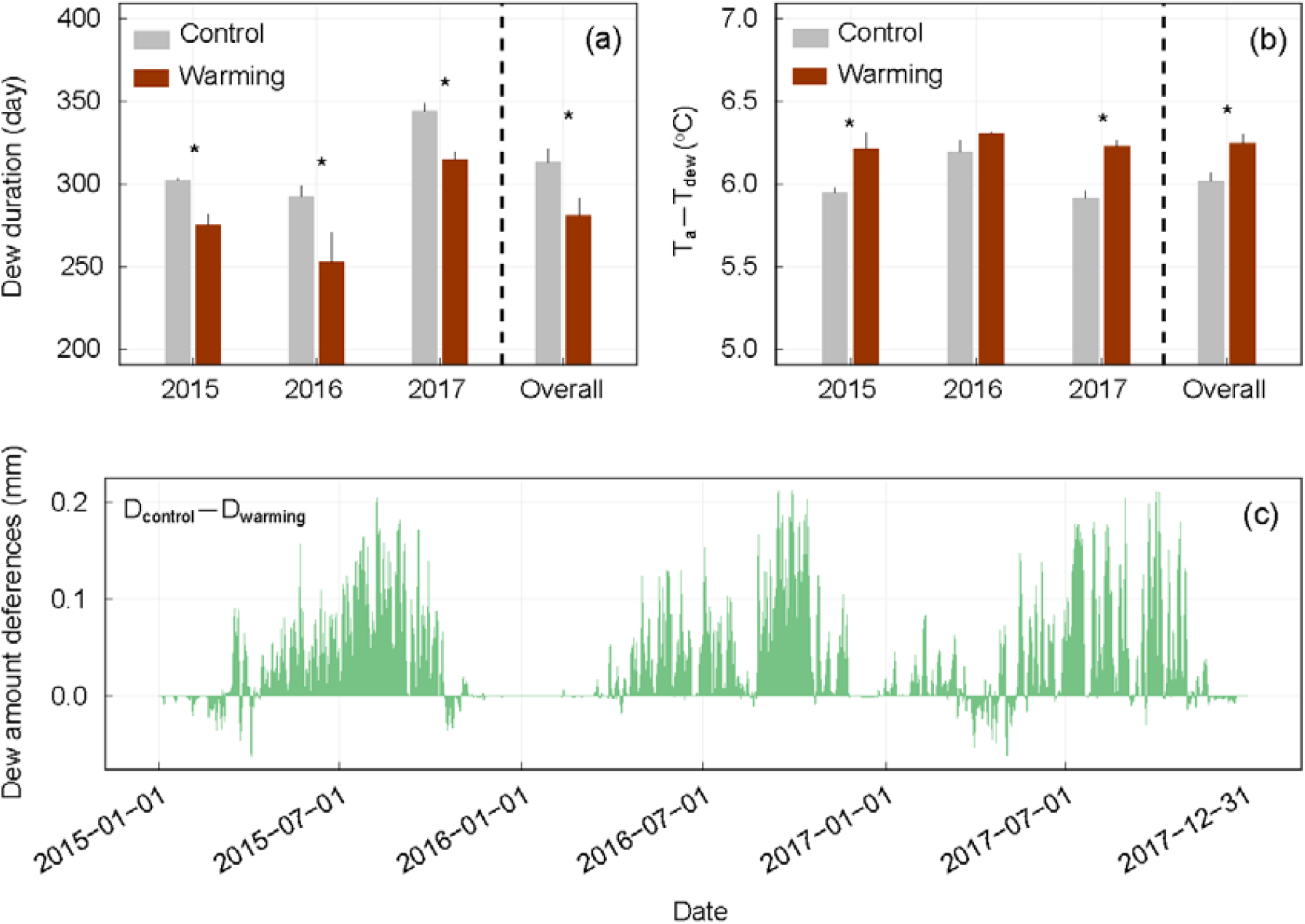
Warming effects on (a) dew duration, (b) the difference between the air temperature (T_a_) and dew point temperature (T_dew_) and (c) the differences of dew amount between the control and warming treatment. * indicates statistically significant at *P* < 0.001.

### 3.2. Effects of warming on dew amount among different functional groups

The total aboveground biomass and dew amounts among each functional group were measured by *in situ* dew formation measurements on plants in this study. The results showed that different plant functional groups significantly differed in dew formation and warming significantly decreased the dew amount among each functional group (a reduction of 83.5%, 71.6%, 97.6% and 87.0% for sedges, forbs, grasses and all species combined, Fig. 3a), while it slightly changed the aboveground biomass of different functional groups (Fig. 3b).

**Fig. 3.**
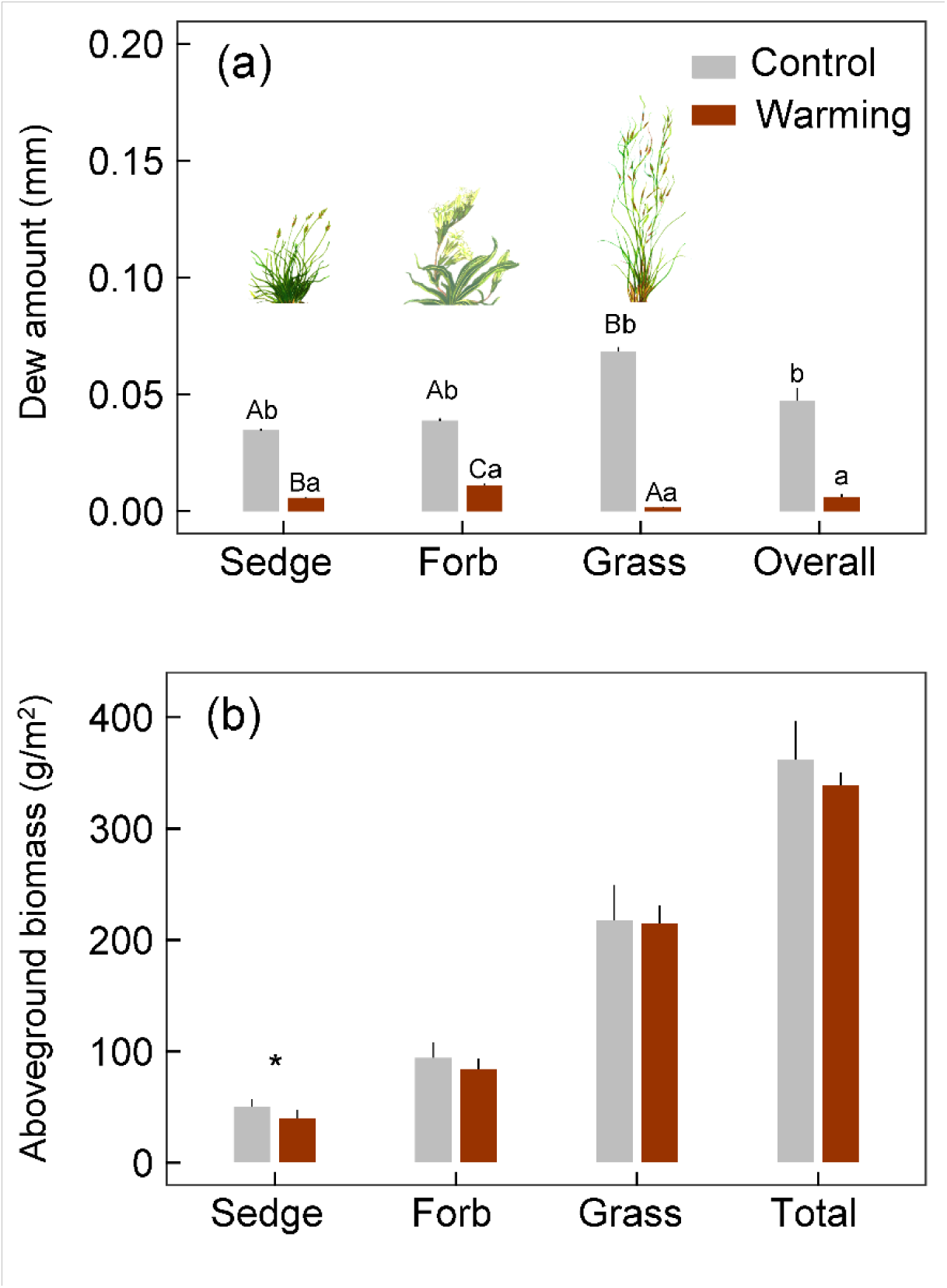
Warming effects on (a) dew amount and (b) aboveground biomass of different functional groups. Different uppercase letters indicate significant difference in different functional groups (*P*<0.05) and different lowercase letters indicate significant difference in control and warming treatments (*P*<0.05).

### 3.3. Effects of warming on the relationships between plant height and dew amount

Compared with the control treatment, the warming treatment significantly affected the relationship between plant height and dew amount (*P* < 0.001, n=60; Fig. 4). In the control treatment, linear regression revealed that the dew amount was significantly positively correlated with plant height (R^2^ = 0.35, *P* < 0.001; Fig. 4a). However, dew amount was significantly negatively correlated with plant height (R^2^ = 0.34, *P* < 0.001; Fig. 4b) in the warming treatment.

**Fig. 4.**
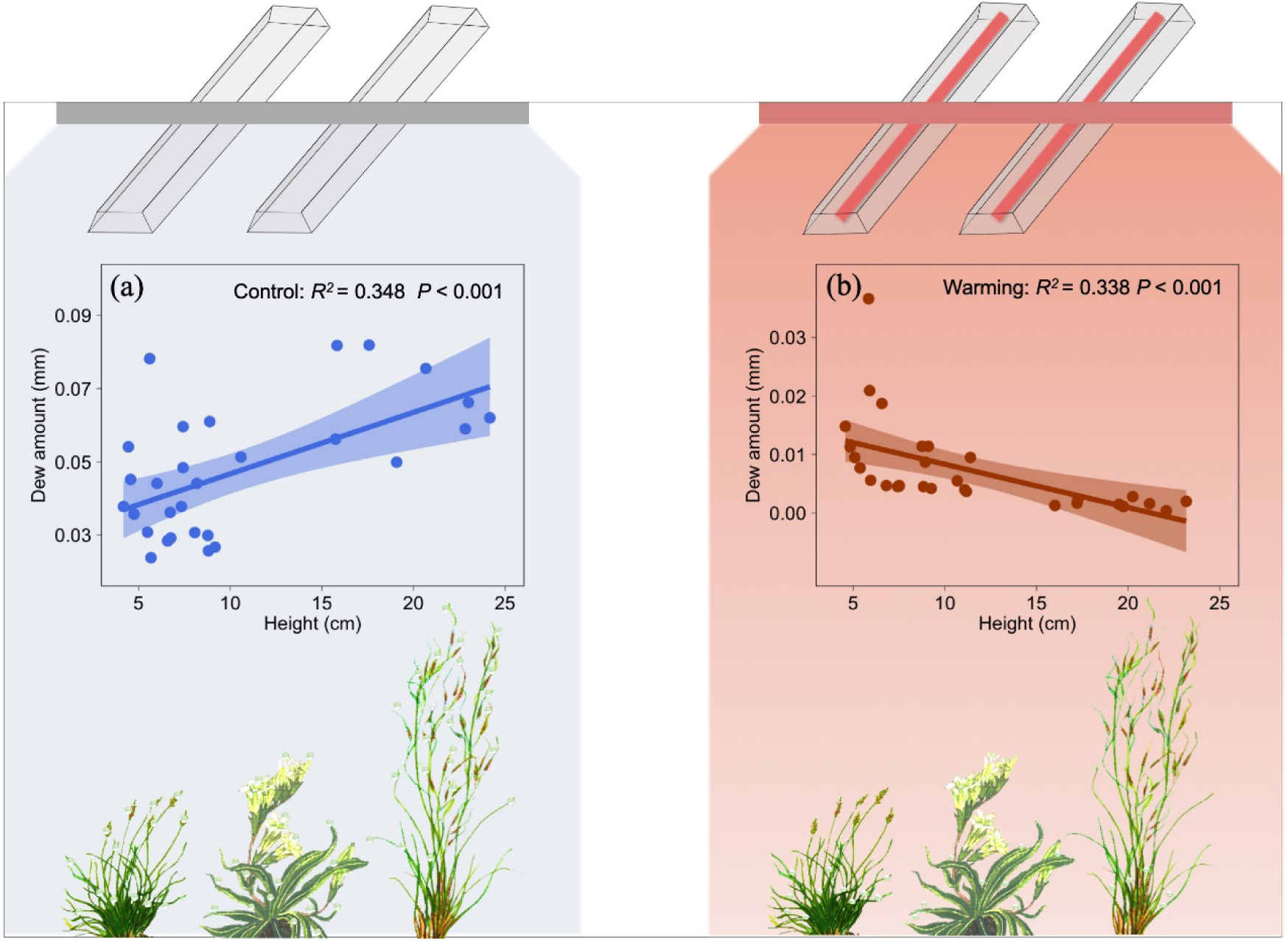
The relationships between plant height and dew amount in control and warming plots.

## 4. Discussion

### 4.1. Warming reduces dew amount and changes seasonal patterns of dew formation

Using three distinct measurement methods, our study showed that warming significantly reduces dew amount (Fig. 1), which may have substantial impacts on plant growth especially during the dry period in the alpine and dryland ecosystems (Stone, 1957; Jacobs et al., 2006; Benasher et al., 2010; Rodney et al., 2013). Warming can reduce dew formation in two ways: by hindering dew condensation and shortening dew retention. Warming can hinder the dew condensation processes by decreasing the air humidity and increasing evaporation (Oliveira 2013; Scheff and Frierson, 2014; Li et al., 2018). Additionally, warming changes the air temperature, dew point temperature and dew point depression (Fig. 2b), which makes it more difficult for the air temperature to approach the dew point temperature (Beysens, 1995; Jacobs et al., 2006; Mortuza et al., 2014). Warming can also accelerate the dew evaporation process (Xiao et al, 2013). Dew droplets lasted for a shorter period of time under warmer temperatures, which also led to a lower dew duration or amount (Xu et al., 2015).

In this study, we found that warming not only reduces dew formation but also changes its seasonal variation (Fig. 2). Therefore, plants growing under water stress would have higher risks of not surviving under warming conditions (Rodney et al., 2013; Tomaszkiewicz et al., 2017). Overall, under the rapidly changing climate, changes in dew formation should be considered an important environmental factor and should not be neglected in arid and cold regions.

### 4.2. Dew formation varied among different functional groups under warming

Functional groups create different microenvironments and have different water use strategies to influence the dew production and absorption by plants (Zhuang and Zhao, 2017; Wang et al., 2019). Environmental conditions, such as temperature, relative humidity and wind speed, change due to various micromorphological features and distribution patterns among different functional groups (Agam and Berliner 2006), affecting dew formation and duration (Ninari and Berliner 2002). Our results showed that different functional groups had different degrees of dew formation, consistent with our expectations.

To date, few studies have investigated how biotic factors (e.g., plant traits and functional groups) affect dew formation. Here, we examined the effects of plant traits (i.e., plant height and aboveground biomass) on dew formation in different plant functional groups (sedges, forbs and grasses) and found that sedges and forbs with shorter heights are associated with less dew than grasses with taller heights under natural conditions (Fig. 3). Because under ambient conditions, the upper canopy air temperature is lower at night due to this area receiving less land-surface radiation, dew formation occurs earlier in higher leaves, such as those of grasses (Zhang et al., 2009; Wang et al., 2017a). In addition, the dominant taller species (*Stipa aliena, Elymus nutans*, and *Helictotrichon tibeticum*) usually have more aboveground biomass (Konrad et al., 2015; Ma et al., 2017) than shorter species, which can facilitate dew formation and retention (Pan et al., 2010). Additionally, the dew water stored within a dense canopy can be preserved for a longer period of time through the reduction in evaporation (Xiao et al., 2013).

Under warming conditions, the aboveground biomass and plant height increased, and the community composition changed with a higher prevalence of grass in the alpine ecosystems (Liu et al., 2018). Such changes should be beneficial for dew formation based on our findings under ambient conditions (i.e., results from the control plots, Fig. 4a). However, a substantial reduction in dew formation was observed under the warming treatments (Fig. 1 and Fig. 2). In addition, we found that warming resulted in a lower dew amount on taller plants, in contrast to the results under ambient conditions (Fig. 4). Warming changed the relationship between plant height and dew amount in both direct and indirect ways. Warming directly affected the air temperature profile and made dew formation more difficult (Wolkovich et al., 2012). In this case, the taller plants had less dew formation because artificial infrared heating made the temperature of the taller canopy higher than that of the lower canopy (Xiao et al., 2013). Warming indirectly caused the soil moisture to evaporate more quickly during the night (Tomaszkiewicz et al., 2015; Li et al., 2018). Therefore, the shorter plants experienced more dew collection than the higher plants during the night under warming conditions. Clearly, warming influenced the dew formation on plants and changed the ecosystem processes compared with those under natural conditions.

### 4.3. Infrared heater warming system reduces dew formation: An overlooked factor in climate change studies

There have been many studies about the response of ecosystem processes to climate change using various artificial warming methods in dry ecosystems (Kimball et al., 2018; Song et al., 2019; Korell et al., 2019), but the possible impacts from the differences between artificial and natural warming on the experimental results have often been overlooked. Our results showed that artificial warming (with an infrared heater warming system) affects dew formation, which likely affects ecosystem processes (Liu et al., 2016). However, it is worth noting that natural climate warming and the infrared heater warming system differ in terms of their heat-dissipating pathways (Korell et al., 2019). Artificial warming generates more heat radiation in the air and drier micro-environments than natural warming (Liu et al., 2018). This difference will affect a number of ecosystem processes and is often overlooked across simulated climate change experiments. Warming makes plants grow taller (Liu et al., 2018), but taller plants produced less dew under warming in our study (Fig. 3). This indicates that the dew formation was significantly reduced under the experimental warming conditions. In addition, the relationship between dew formation and plant height changed being positively correlated under the control treatment to being negatively correlated under the warming treatment (Fig. 4). For such cases, the conclusions of the impacts of warming obtained by artificial warming experiments may deviate from the actual impacts of warming on ecosystem processes. Under future climate warming, the changes in water condensation will also have an especially profound impact on the ecosystem patterns and processes in dryland ecosystems (Li et al., 2018; Wang et al., 2017b). Therefore, we suggest that the impact of experimental warming on dew formation should be considered an important environmental factor affecting ecosystems processes during climate warming.

## 5. Conclusions

Using three measurement methods, we observed that warming significantly reduced the dew amount and duration and changed its seasonal patterns. Different plant functional groups had different effects on dew formation due to their associated microclimates and plant heights, resulting in taller plants experiencing more dew formation. However, artificial warming caused the taller plants to have less dew formation due to the associated heat radiation. We also found that infrared heater warming systems markedly reduced dew formation, which should be addressed to avoid overestimating the impact of climate warming on ecosystems during global change studies. Our study demonstrates that dew condensation responds to climate warming and highlights that microhabitat conditions and plant traits mediate dew formation under warming conditions, having an important potential effect on ecosystems processes in the future.

## Authors’ contributions

Jin-sheng He conceived the ideas and designed methodology; Lixu Zhang and Qian Chen collected the data; Lixu Zhang, Zhiyuan Ma, Zijian Shangguan, Hao Wang and Tianjiao Feng analysed the data; Lixu Zhang and Tianjiao Feng led the writing of the manuscript with the assistances from Lixin Wang. All authors contributed critically to the drafts and gave final approval for publication.

## Acknowledgments

This research was funded by the National Natural Science Foundation of China (31630009, 31621091), and the 111 Project (Grant No. B14001). We sincerely thank the contribution and assistance of Haibei National Field Research Station of Alpine Grassland Ecosystem.

---

**Fig. S1.**
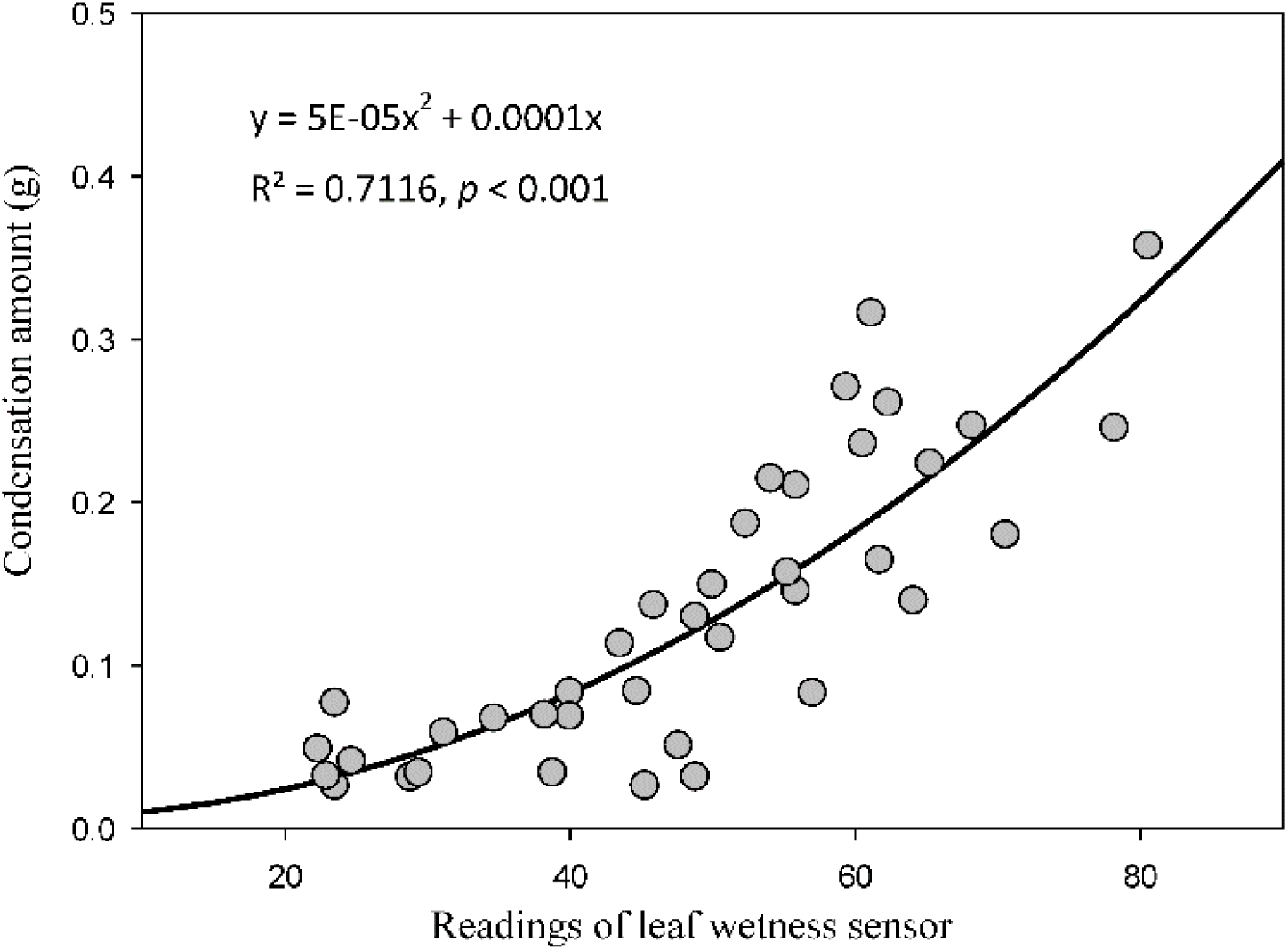
Fitting curve of the readings of leaf wetness sensor and condensation amount.

## References

Agam, N., and P. R. Berliner. 2006. Dew formation and water vapor adsorption in semi-arid environments--A review. Journal of Arid Environments 65:572–590.

Allen, R. G., L. S. Pereira, D. Raes, and Smith M. 1998. Crop evapotranspiration: guidelines for computing crop water requirements. Food and Agriculture Organization of the United Nations, Rome.

Benasher, J., P. Alpert, and A. Benzvi. 2010. Dew is a major factor affecting vegetation water use efficiency rather than a source of water in the eastern Mediterranean area. Water Resources Research 46:437–441.

Beysens, D. 1995. The formation of dew. Atmospheric Research 39:215–237.

Beysens, D. 2016. Estimating dew yield worldwide from a few meteo data. Atmospheric Research 167:146–155.

Chen, Q. 2015. Hydrological processes and their response to warming and altered precipitation in an alpine meadow on the Tibetan Plateau (Master’s thesis). Peking University, Beijing.

De Boeck, H. J., S. Vicca, J. Roy, I. Nijs, A. Milcu, J. Kreyling, A. Jentsch, A. Chabbi, M. Campioli, T. Callaghan, C. Beierkuhnlein, and C. Beier. 2015. Global Change Experiments: Challenges and Opportunities. BioScience 65:922–931.

Ettinger, A. K., I. Chuine, B. I. Cook, J. S. Dukes, A. M. Ellison, M. R. Johnston, A. M. Panetta, C. R. Rollinson, Y. Vitasse, E. M. Wolkovich, and E. Jeffers. 2019. How do climate change experiments alter plot-scale climate? Ecology Letters 22:748–763.

Fischer, T., M. Veste, O. Bens, and R. F. Huttl. 2012. Dew formation on the surface of biological soil crusts in central European sand ecosystems. Biogeosciences 9:4621–4628.

Goldsmith, G. R., N. J. Matzke, and T. E. Dawson. 2013. The incidence and implications of clouds for cloud forest plant water relations. Ecology Letters 16:307–314.

Hao, X., C. Li, B. Guo, J. Ma, M. Ayup, and Z, Chen. 2012. Dew formation and its long-term trend in a desert riparian forest ecosystem on the eastern edge of the Taklimakan Desert in China. Journal of Hydrology 472-473:90–98.

Jacobs, A. F. G., B. G. Heusinkveld, R. J. W. Kruit, and S. M. Berkowicz. 2006. Contribution of dew to the water budget of a grassland area in the Netherlands. Water Resources Research 42:446–455.

Kidron, G. J., and M. Temina. 2013. The Effect of Dew and Fog on Lithic Lichens Along an Altitudinal Gradient in the Negev Desert. Geomicrobiology Journal 30:281–290.

Kimball, B. A., A. M. Alonso-Rodriguez, M. A. Cavaleri, S. C. Reed, G. Gonzalez, and T. E. Wood. 2018. Infrared heater system for warming tropical forest understory plants and soils. Ecoloy Evolution 8:1932–1944.

Konrad, W., J. Burkhardt, M. Ebner, and A. Roth-Nebelsick. 2015. Leaf pubescence as a possibility to increase water use efficiency by promoting condensation. Ecohydrology 8:480–492.

Korell, L., H. Auge, J. M. Chase, W. S. Harpole, and T. M. Knight. 2019. We need more realistic climate change experiments for understanding ecosystems of the future. Globe Change Biology. doi: 10.1111/gcb.14797.

Li, X. R., R. L. Jia, Z. S. Zhang, P. Zhang, and R. Hui. 2018. Hydrological response of biological soil crusts to global warming: A ten-year simulative study. Global Change Biology 24:4960–4971.

Lin, L., B. Zhu, C. Chen, Z. Zhang, Q. B. Wang, and J. S. He. 2016. Precipitation overrides warming in mediating soil nitrogen pools in an alpine grassland ecosystem on the Tibetan Plateau. Scitific Reports 6:31438.

Liu, H. Y., Z. R. Mi, L. Lin, Y. H. Wang, Z. H. Zhang, F. W. Zhang, H. Wang, L. L. Liu, B. A. Zhu, G. M. Cao, X. Q. Zhao, N. J. Sanders, A. T. Classen, P. B. Reich, and J. S. He. 2018. Shifting plant species composition in response to climate change stabilizes grassland primary production. Proceedings of the National Academy of Sciences of the United States of America 115:4051–4056.

Liu, L. L., X. Wang, M. J. Lajeunesse, G. F. Miao, S. L. Piao, S. Q. Wan, Y. X. Wu, Z. H. Wang, S. Yang, P. Li, and M. F. Deng. 2016. A cross-biome synthesis of soil respiration and its determinants under simulated precipitation changes. Global Change Biology 22:1394–1405.

Ma, Z., H. Liu, Z. Mi, Z. Zhang, Y. Wang, W. Xu, L. Jiang, and J. S. He. 2017. Climate warming reduces the temporal stability of plant community biomass production. Nature Communication 8:15378.

Moni, C., H. Silvennoinen, B. A. Kimball, E. Fjelldal, M. Brenden, I. Burud, A. Flo, and D. P. Rasse. 2019. Controlled infrared heating of an artic meadow: challenge in the vegetation establishment stage. Plant Methods 15:1–12.

Mortuza, M. R., S. Selmi, M. M. Khudri, A. K. Ankur, and M. M. Rahman. 2014. Evaluation of temporal and spatial trends in relative humidity and dew point temperature in Bangladesh. Arabian Journal of Geosciences 7:5037–5050.

Munné, S. 1999. Role of Dew on the Recovery of Water-Stressed Melissa officinalis L. Plants. Animal Behaviour 72:1405–1416.

Ninari, N., and P. R. Berliner. 2002. The role of dew in the water and heat balance of bare loess soil in the Negev Desert: quantifying the actual dew deposition on the soil surface. Atmospheric Research 64:323–334.

Oliveira, R. S. 2013. Foliar uptake of fog water and transport belowground alleviates drought effects in the cloud forest tree species, Drimys brasiliensis (Winteraceae). New Phytologist 199:151–162.

Pan, Y. X., X. P. Wang, and Y. F. Zhang. 2010. Dew formation characteristics in a revegetation-stabilized desert ecosystem in Shapotou area, Northern China. Journal of Hydrology 387:265–272.

Rodney, E. W., M. W. Stuart, B. Z. Chris, and C. H. Thomas. 2013. Increased vapor pressure deficit due to higher temperature leads to greater transpiration and faster mortality during drought for tree seedlings common to the forest-grassland ecotone. The New Phytologist 200:366–374.

Scheff, J., and D. M. W. Frierson. 2014. Scaling Potential Evapotranspiration with Greenhouse Warming. Journal of Climate 27:1539–1558.

Shaver GR, Canadell J, Chapin FS III, Gurevitch J, Harte J, Henry G, Ineson P, Jonasson S, Melillo J, Pitelka L, Rustad L. 2000. Climate warming and terrestrial ecosystems: a conceptual framework for analysis. Bioscience 50: 871–882.

Song, J., S. Wan, S. Piao, A. K. Knapp, A. T. Classen, S. Vicca, P. Ciais, M. J. Hovenden, S. Leuzinger, C. Beier, P. Kardol, J. Y. Xia, Q. Liu, J. Y. Ru, Z. X. Zhou, Y. Q. Luo, D. L. Guo, J. A. Langley, J. Zscheischler, J. S. Dukes, J. W. Tang, J. Q. Chen, K. S. Hofmockel, L. M. Kueppers, L. Rustad, L. L. Liu, M. D. Smith, P. H. Templer, R. Q. Thomas, R. J. Norby, R. P. Phillips, S. L. Niu, S. Fatichi, Y. P. Wang, P. S. Shao, H. Y. Han, D. D. Wang, L. J. Lei, J. L. Wang, X. N. Li, Q. Zhang, X. M. Li, F. L. Su, B. Liu, F. Yang, G. G. Ma, G. Y. Li, Y. C. Liu, Y. Z. Liu, Z. L. Yang, K. S. Zhang, Y. Miao, M. J. Hu, C. Yan, A. Zhang, M. X. Zhong, Y. Hui, Y. Li, and M. M. Zheng. 2019. A meta-analysis of 1,119 manipulative experiments on terrestrial carbon-cycling responses to global change. Nature Ecology & Evolution 3:1309–1320.

Steinberger, Y., I. Loboda, and W. Garner. 1989. The influence of autumn dewfall on spatial and temporal distribution of nematodes in the desert ecosystem. Journal of Arid Environments 16:177–183.

Stone, E. C. 1957. Dew as an Ecological Factor: I. A Review of the Literature. Ecology 38:407–413.

Tomaszkiewicz, M., M. Abou Najm, D. Beysens, I. Alameddine, and M. El-Fadel. 2015. Dew as a sustainable non-conventional water resource: a critical review. Environmental Reviews 23:425–442.

Tomaszkiewicz, M., M. A. Najm, R. Zurayk, and M. El-Fadel. 2017. Dew as an adaptation measure to meet water demand in agriculture and reforestation. Agricultural & Forest Meteorology 232:411–421.

Tomaszkiewicz, M., M. Abou Najm, D. Beysens, I. Alameddine, E. B. Zeid, and M. El-Fadel. 2016. Projected climate change impacts upon dew yield in the Mediterranean basin. Science of the Total Environment 566:1339–1348.

Vuollekoski, H., M. Vogt, V. A. Sinclair, J. Duplissy, H. Järvinen, E. M. Kyrö, R. Makkonen, T. Petäjä, N. L. Prisle, and P. Räisänen. 2015. Estimates of global dew collection potential on artificial surfaces. Hydrology & Earth System Sciences 11:601–613.

Walther, G. R., E. Post, P. Convey, A. Menzel, C. Parmesan, T. J. C. Beebee, J. M. Fromentin, O. Hoegh-Guldberg, and F. Bairlein. 2002. Ecological responses to recent climate change. Nature 416:389–395.

Wang, L., K. F. Kaseke, and M. K. Seely. 2017a. Effects of non-rainfall water inputs on ecosystem functions. Wiley Interdisciplinary Reviews: Water 4:e1179.

Wang, L., K. F. Kaseke, S. Ravi, W. Jiao, R. Mushi, T. Shuuya, and G. Maggs-Kölling. 2019. Convergent vegetation fog and dew water use in the Namib Desert. Ecohydrology. DOI: 10.1002/eco.2130.

Wang, J., L. Liu, X. Wang, S. Yang, B. Zhang, P. Li, C. Qiao, M. Deng, W. Liu, and K. Treseder. 2017b. High night-time humidity and dissolved organic carbon content support rapid decomposition of standing litter in a semi-arid landscape. Functional Ecology 31:1659–1668.

Wang, G., G. Liu, and C. Li. 2012. Effects of changes in alpine grassland vegetation cover on hillslope hydrological processes in a permafrost watershed. Journal of Hydrology 444:22–33.

Wolkovich, E. M., B. I. Cook, J. M. Allen, T. M. Crimmins, J. L. Betancourt, S. E. Travers, S. Pau, J. Regetz, T. J. Davies, N. J. B. Kraft, T. R. Ault, K. Bolmgren, S. J. Mazer, G. J. McCabe, B. J. McGill, C. Parmesan, N. Salamin, M. D. Schwartz, and E. E. Cleland. 2012. Warming experiments underpredict plant phenological responses to climate change. Nature 485:494–497.

Xiao, H., R. Meissner, J. Seeger, H. Rupp, H. Borg, and Y. Zhang. 2013. Analysis of the effect of meteorological factors on dewfall. Science of the Total Environment 452-453:384–393.

Xu, Y., B. Yan, and J. Tang. 2015. The Effect of Climate Change on Variations in Dew Amount in a Paddy Ecosystem of the Sanjiang Plain, China. Advances in Meteorology 9:1–9.

Zhang, J., Y. M. Zhang, A. Downing, J. H. Cheng, X. B. Zhou, and B. C. Zhang. 2009. The influence of biological soil crusts on dew deposition in Gurbantunggut Desert, Northwestern China. Journal of Hydrology 379:220–228.

Zheng, Y., H. Bai, Z. Huang, X. Tian, F. Q. Nie, Y. Zhao, J. Zhai, and L. Jiang. 2010. Directional water collection on wetted spider silk. Nature 463:640–643.

Zhuang, Y., and S. Ratcliffe. 2012. Relationship between dew presence and Bassia dasyphylla plant growth. Journal of Arid Land 4:11–18.

Zhuang, Y. L., and W. Z. Zhao. 2017. Dew formation and its variation in Haloxylon ammodendron plantations at the edge of a desert oasis, northwestern China. Agricultural and Forest Meteorology 247:541–550.

